# Gut microbiome, diet and symptom interactions in irritable bowel syndrome

**DOI:** 10.1101/2020.02.25.964528

**Authors:** Julien Tap, Stine Störsrud, Boris Le Nevé, Aurélie Cotillard, Nicolas Pons, Joël Doré, Lena Öhman, Hans Törnblom, Muriel Derrien, Magnus Simren

## Abstract

While several studies have documented associations between dietary habits and microbiota composition and function in healthy subjects, no study explored these associations in patients with irritable bowel syndrome (IBS), and especially in relation to symptoms. Here, we used a novel approach that combined data from 4-day food diary, integrated into a food tree, together with gut microbiota (shotgun metagenomic) for IBS patients (*N*=149) and healthy subjects (*N*=52). Paired microbiota and food-based trees allowed to detect new association between subspecies and diet. Combining co-inertia analysis and linear regression models, exhaled gas levels and symptom severity could be predicted from metagenomic and dietary data. IBS patients with severe symptoms had a diet enriched in food items of poorer quality, a high abundance of gut microbial enzymes involved in hydrogen metabolism in correlation with animal carbohydrate (mucin/meat-derived) metabolism. Our study provides unprecedented resolution of diet-microbiota-symptom interactions and ultimately paves the way for personalized nutritional recommendations.

## Introduction

Irritable bowel syndrome (IBS) is one of the most common gastrointestinal disorders, affecting approximately 10 % of the population ^1^. However, the effectiveness of treatment options for this common disorder is limited, and IBS is associated with profound reduction in health-related quality of life, and huge societal costs ^2^. Most patients with IBS report the triggering or worsening of symptoms following food intake ^3^. Therefore, the interest in the dietary management of IBS symptoms has increased, particularly over the last decade. Recent clinical studies have shed light on the importance of specific food items in the exacerbation of gastrointestinal (GI) symptoms. However, the optimal dietary recommendations for individual IBS patients still remains uncertain ^3^. Diet, particularly long-term eating habits, is known to be one of the drivers of microbiota variation. Detailed tracking of the covariation between the gut microbiota and diet has opened up new perspectives for personalization of the gut microbiota response to diet ^4, 5^.

In population-based cohorts, significant associations have been found between dietary factors and interindividual distances in microbiota composition ^6, 7, 8, 9, 10^. Culture-based studies have shown that strains from a given species may share substrate metabolism but may also display differences ^11, 12^. Consistent with this finding, recent metagenomics-based studies using strain-level profiling tools have revealed functional variation within species associated with dietary habits ^13, 14, 15^.

Carbohydrates are among the food items that can exacerbate symptoms, particularly those that are incompletely absorbed in the small intestine, which lead to gas accumulation including hydrogen and methane when microbial fermentation occurs in the colon ^16, 17^. These carbohydrates include among others short-chain fermentable carbohydrates (fermentable oligosaccharide, disaccharide, monosaccharide and polyol (FODMAP) ^18^ which restriction has been associated with alteration in gut microbiota ^16^. The gut microbiota has a far greater capacity to encode enzymes for metabolizing food glycans, (CAZy, or carbohydrate-active enzymes) than the human genome ^19^, and CAZy profiles have been associated with different dietary habits ^20, 21^, and host parameters ^22^.

We previously showed that the gut microbiota is associated with symptom severity in IBS, and identified a microbial signature for IBS severity that was not associated with intake of macronutrients ^23^. Although considered to be associated with symptom generation in patients with IBS, the way in which the combination of diet and gut microbiota affects symptoms remains unknown ^23, 24, 25^. Therefore, in this study, we investigated the relationship between digestive symptoms and extensive datasets of diet and the gut microbiota of IBS patients, with a focus on IBS patients with severe symptoms, further referred as severe IBS, making use of a whole-metagenomics sequencing approach and categorized dietary intake based on a 4-day food diary.

First, using a nutritional-based diet index, we showed that IBS patients with severe symptoms, are characterized by a higher intake of food items of poorer quality during their main meals. Then, combining co-inertia analysis and linear regression analysis suggested that gut microbial hydrogen metabolism and dietary profile are associated with IBS symptom severity. Additionally, our study further suggests that specific hydrogenases were associated with gut microbiota function in terms of the CAZy involved in animal carbohydrate metabolism.

## Results

### A food item-based tree differentiates between the dietary habits of IBS patients and healthy subjects

We investigated the association of GI symptom severity and patterns with diet and the gut metagenome in 52 healthy subjects and 149 IBS patients (Table 1). The IBS symptoms were severe, according to the IBS Severity Scoring System (IBS-SSS) ^26^, in 65 of the 149 IBS patients. A 4-day food diary was obtained from 142 subjects. Dietary macronutrients and micronutrients identified as consumed differentially between IBS patients and healthy subjects or between patients with severe IBS symptoms and other individuals (healthy or with non-severe, i.e. mild/moderate IBS symptoms) did not remain significant after correction for multiple testing (Fig. S1 and S2). This suggests that dietary data, aggregated at the level of micro- and macronutrients, do not differ between IBS patients and healthy subjects, and neither along severity gradient.

**Table 1:**
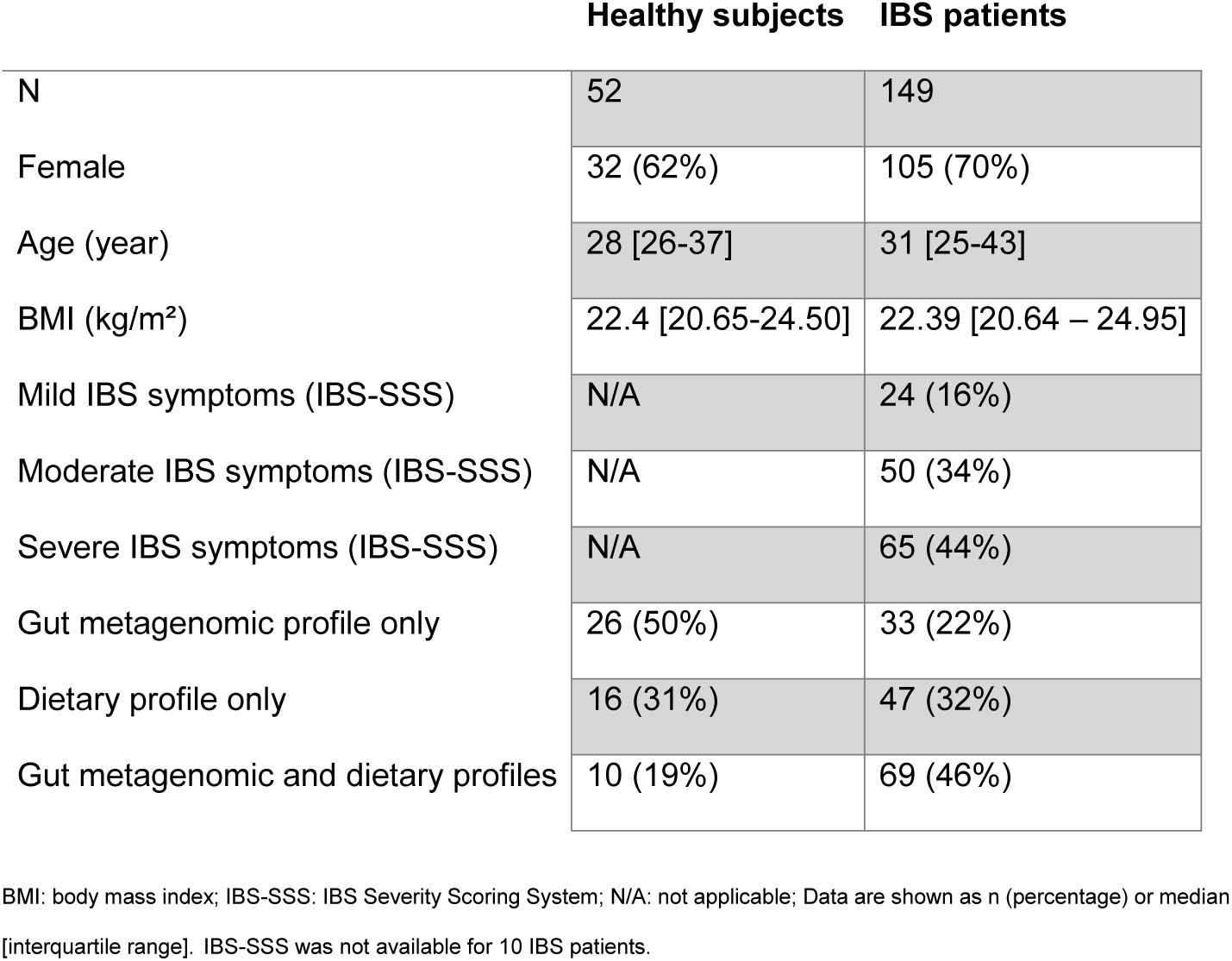
Characteristics of the study cohort.

Next, we assembled a food item tree, similar to that used by Johnson et al. ^4^, clustering food items on the basis of nutritional content rather than the food item itself. In total, the participants consumed 966 different food items, which were aggregated into three hierarchical levels, based on the National Swedish food database, to build a hierarchical tree. A fourth level based on nutrient composition was also added, resulting in 85 food-nutrient groups (Fig. 1a and Table S1). The unweighted UniFrac distance between individuals was calculated using the four hierarchical levels of dietary data and subjected to principal coordinate analysis (PCoA) (Fig. 1b). Based on Spearman’s correlation coefficient, the four hierarchical food levels, including food-nutrient groups, could be projected onto principal coordinate axes 1 and 2 (PCoA1 and PCoA2). PCoA1 (7.9% of total variance) was associated with the separation of subjects according to their consumption of meat-based and plant-based food items (Fig. 1b). Further, PCoA2 (6.1% of total variance) separated subjects on the basis of their consumption of unprocessed food items, such as fish and eggs, and processed food items, such as candy and fried potato products. Following the integration of selected clinical variables into the analysis, PCoA1 was found to be associated with sex (Fig. 1c, Mann-Whitney test, p<0.05) whereas PCoA2 tended to be associated with IBS severity (Fig. 1d, Mann-Whitney test, p=0.06). A meat-to-plant ratio was calculated for each subject, based on the aggregation of food items into these two categories (Table S2). PCoA1 was significantly associated with the meat/plant ratio (Fig. 1e, rho = −0.62, p<0.05), suggesting that the proportion of meat-based food relative to plant-based food was the major driver of dietary variation between subjects in our cohort.

**Figure 1:**
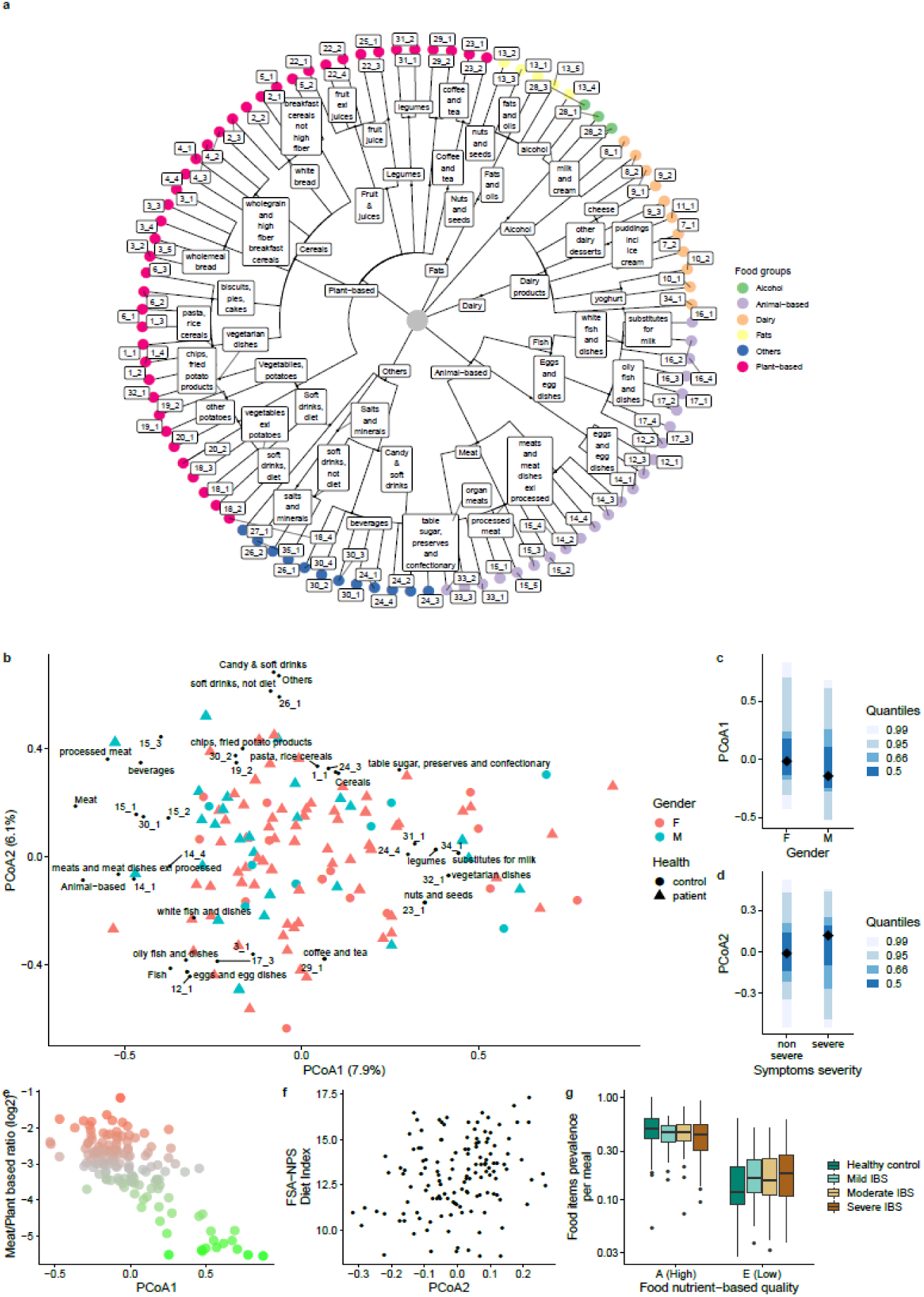
Quantity and quality assessment for dietary profiles and analyses of associations with gastrointestinal symptom severity. **a** Food item-based hierarchical tree based on the National Swedish food database and nutrient based clustering. **b** Principal coordinate analysis of unweighted UniFrac distance between the dietary profiles of individuals. Food levels were projected onto the two first coordinates (PCoA1 and PCoA2) based on Spearman’s correlation analyses (see Supp Table 1 for terminology of food level 4). Color indicates the sex of the individual and the shape of the point indicates health status. **c** PCoA1 as a function of sex. **d** PCoA2 as a function of IBS symptom severity **e** Log2 meat/plant ratio as a function of PCoA1. The color gradient extends from red (all meat) to green (all plant-based foods). **f** FSA-NPS diet index as a function of PCoA2. **g** Prevalence of food items per meal as a function of FSA-NPS food quality and health status. Class A corresponds to high-quality food, whereas class E corresponds to low-quality food.

We then assessed diet quality by performing nutrient profiling according to the Nutrient Profiling System of the British Food Standards Agency (FSA-NSP) for each food item and calculating an individual FSA-NPS Dietary Index (DI) score. A higher FSA-NPS DI reflects a lower nutritional quality of the foods consumed by the individual. The overall dietary quality was not correlated to PCoA1, but was positively correlated with PCoA2, suggesting that IBS symptom severity may be associated with a lower overall dietary quality (Fig. 1f rho = 0.24, p<0.05) although not directly (Mann-Whitney test, p>0.05). With the FSA-NPS, food items could be classified into five groups (from high-quality A, to low-quality E, see methods). IBS patients with severe symptoms consumed higher amount of food items of poorer quality during their main meals than healthy subjects and IBS patients with milder symptoms (Fig. 1g, Mann-Whitney, p<0.05 for high and low food quality). This association was lost when including snacks in the analysis.

Finally, based on the PCoA results, we investigated whether FODMAPs (fermentable oligo-, di-, and monosaccharides and polyols), were associated with diet quality based on the hierarchical food tree. GOS and fructans intakes were positively associated with PCoA1 (rho=0.28 and rho=0.33 respectively, p<0.05), suggesting that subjects who consumed high levels of plant-based food items had a diet enriched in GOS and fructans. Polyol intake was associated with PCoA2 (rho=-0.31, p<0.05), indicating that subjects who consumed high levels of processed foods and lower quality food items had a diet enriched in polyols. The intake of lactose and fructose was not associated with PCoA1 or PCoA2, suggesting that their consumption was not a major driver of global diet variation or symptom generation in our study cohort.

### Dietary profile associations with gut microbiota composition and function

As diet is a major driver of microbiota composition and function, we investigated whether differences previously observed in dietary profile between study subjects, were associated with gut microbiota composition and functions. Enterotypes, previously assessed by 16S rRNA gene sequencing, could be separated into three microbiota communities on the basis of Dirichlet multinomial mixture (DMM) modeling ^23^. Enterotypes were not associated with dietary distance (permanova, p>0.05), nor with meat/plant ratio (p>0.05) or diet quality (FSA-NPS DI, p>0.05), suggesting that dietary variations within this cohort did not have a major effect on global microbiota assemblage.

Next, whole-metagenomic sequencing was performed on 138 individuals (102 IBS patients and 36 healthy subjects) (Table 1). Metagenomic reads (with an average of 14 million reads per sample) were mapped onto a catalog of Metagenomic Species Pangenomes (MSPs) ^27^, yielding a total of 1,661 MSPs. On the basis of per-individual genetic content, 166 of them were further divided into 523 subspecies, corresponding to a mean of 75.3% of the metagenome read mass. The remaining 1495 MSPs were not assigned to MSP subspecies (MSP_unassigned). We then used the taxonomic tree to investigate the effect on dietary profile variation of each taxonomic tree node, from phylum to subspecies level (permanova test). Each microbial lineage with at least one node with an effect size of more than 2% was selected with no filter applied on statistical significance (Fig. 2a). Depending on taxonomic lineage, dietary variation was either better explained at the species (i.e. MSPs) or subspecies level (Fig. 2b). For example, the MSP assigned to *Eubacterium rectale* was less associated with diet than its two subspecies. In particular, the relative abundance of the *E. rectale* subspecies with flagellin-encoding genes (Fig. 2c) correlated with a diet enriched in meat-based products (Rho=0.23, p=0.05) and with vitamin B12 intake (Rho=0.31, p<0.05).

**Figure 2:**
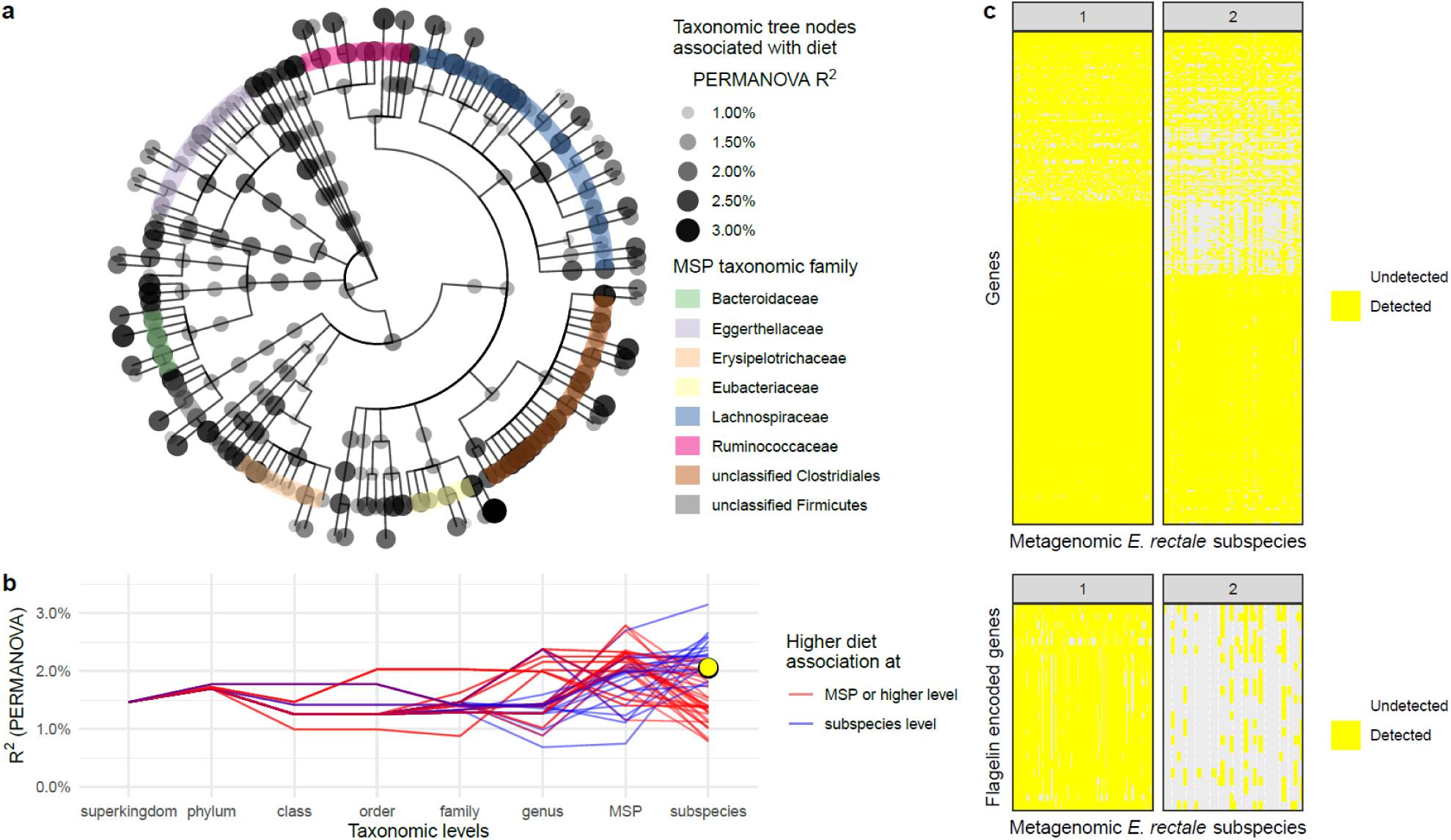
Dietary profile associations with gut microbial lineage and functions. **a** taxonomic tree of the microbial lineage with nodes corresponding to effect sizes of more than 2% (R2 assessed by PERMANOVA). The color code indicates the taxonomic family of the MSP taxonomic family. **b** Effect size of dietary variation (R2 assessed by PERMANOVA) as a function of taxonomic level. Red lines correspond to lineages for which effect size was greater at subspecies level. Yellow dot account for *Eubacterium rectale* subspecies 1. **c** Presence/absence heatmap showing the genes detected within each *Eubacterium rectale* subspecies with a specific focus on flagellin encoded genes. Genes are represented as rows and samples as columns.

### Association of microbiota function and dietary variation with clinical parameters and gas metabolism

As dietary variation was better explained at the species (i.e. MSPs) or subspecies level MSP, we further combined microbiota profile variation (JSD distance at subspecies and MSP_unassigned) with dietary profile variation. Using a co-inertia approach on PCoA components, we explored the complex association between microbiota, diet and multiple explanatory variables in 79 subjects for whom both dietary and microbiota datasets were available (training set, co-inertia RV coefficient=0.59). Subjects with only one of these datasets (microbiota or diet, Table 1) constituted the test set (*N*=122). All 201 study subjects (training and test sets) were projected onto the same hyperspace on the basis of their microbiota and dietary profiles (Fig. 3a). We then investigated how this common projection of gut microbiota and diet was associated with clinical parameters. The relationship of microbiota and dietary profiles to gas metabolism and clinical parameters was investigated by extracting the first co-inertia coordinates from both the training and test sets. Then, we correlated them with variables, including BMI, age, IBS symptoms severity score, exhaled gas metabolism (H_2_ and CH_4_), microbiota gene richness and dietary variables (meat/plant ratio, diet quality) (Fig. 3b). The first seven coordinates (axis A1 to axis A7, Fig. 3c), accounting for 50% of co-variation, were retained. Consistently for both the training and test sets, the meat/plant ratio was associated with axis A1, whereas microbial gene richness, diet quality and exhaled CH_4_ were associated with axis A2 (Table S3). This suggests that meat/plant ratio was the main tested factor explaining microbiota and dietary co-variations independently of gene richness, diet quality and exhaled CH_4_. Although weaker, consistent correlation directions for axis A1, in both the test and training set, were detected for exhaled H_2_ and IBS symptom severity. Since CH_4_ metabolism is dependent on H_2_ metabolism, the ratio of exhaled H_2_ to exhaled CH_4_ for each subject was calculated. This ratio was consistently correlated with axis 4, in both the food and microbiota in the training sets and microbiota in test set, suggesting that H_2_/CH_4_ metabolism can be explained by metagenomics and dietary profile variations.

**Figure 3:**
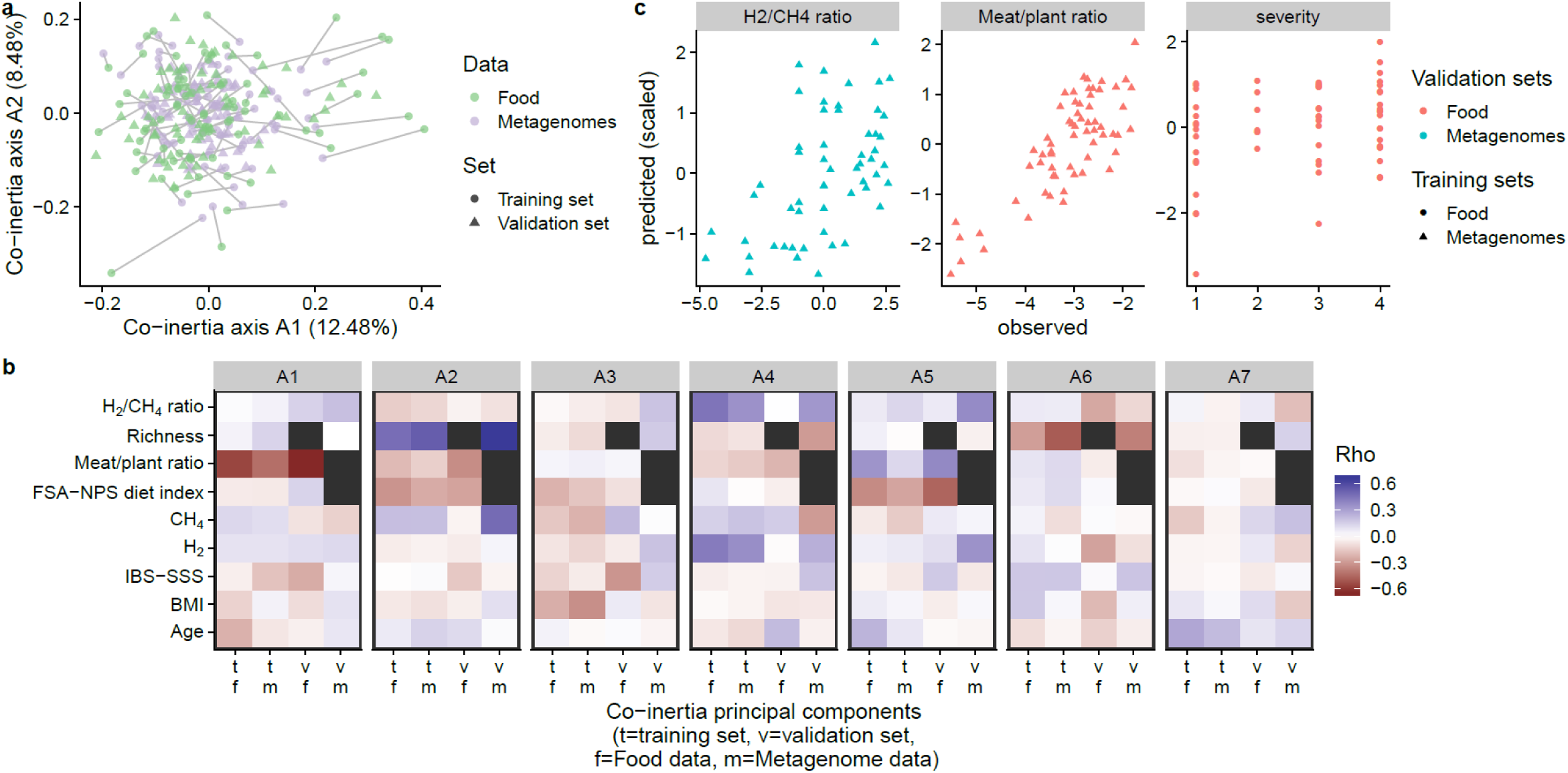
Co-inertia analysis associates microbiota profiles with dietary profiles, together with gas metabolism and symptom severity. **a** Co-inertia scatter plot with the training set (*n*=79) including both metagenomic and dietary data, and the test set (*n*=122) for which only dietary or metagenomics data were available. Individual coordinates for the test set were computed from their PCoA coordinates with the co-inertia model. Color indicates the data source (diet or metagenomic). **b** Heatmap of Spearman’s correlations between the first seven co-inertia axes and clinical, gas metabolism, gene richness and dietary data. The color indicates the strength of the correlation. Black squares indicate missing data. v: validation set; t: training set; m: microbiota; f: food. **c** Scatter plots of the relationship between predicted (linear regression) and observed data for H_2_/CH_4_ ratio, meat/plant ratio and symptom severity (based on the first 5 co-inertia components). The color indicates the validation set source, whereas the shape of the points indicates the training set source (dietary or metagenomic). IBS symptom severity groups (healthy, mild, moderate and severe) were coded from 1 to 4, respectively.

For the prediction of each clinical variable, we used a machine learning approach by constructing regression models from the first five co-inertia coordinates (40% of the variance) extracted from the microbiota and dietary datasets. Co-inertia models trained on microbiota and validated on dietary data were assessed for robustness by regression analysis (Fig. 3c). For example, using a model fitted on the microbiota training set, H_2_/CH_4_ ratio could be predicted from the microbiota test set (Pearson r=0.53, *p*<0.05). Similarly, using a model fitted on the microbiota training set, meat/plant ratio could be predicted from the dietary test set (r=0.69, *p*<0.05). Using another model fitted on the dietary training set, IBS symptom severity could be predicted from the dietary test set (r=0.28, *p*<0.05).

### Carbohydrate-associated enzymes and hydrogenases encoded by metagenomes are associated with IBS symptom severity

As carbohydrates are among the food items that can exacerbate symptoms which lead to gas accumulation, we further explored the relation between CAZy and hydrogenases. Metagenomes were clustered according to DMM models based on the relative abundance of CAZy. Using the minimum Laplace approximation, study subjects could be divided into three distinct CAZy clusters (CAZotype), as enterotypes, which were significantly associated (chi^2^, *p*<0.05, Fig. S4), suggesting a link between carbohydrate metabolism and enterotypes. In contrast to enterotypes, CAZotypes were associated with diet profile variation (permanova, p<0.01).

We investigated potential links between CAZy and hydrogenases encoded by gut metagenomes and symptom severity. CAZy were subdivided into broad substrate categories (plant cell-wall carbohydrates, animal carbohydrates, peptidoglycan, and others (starch/glycogen, sucrose/fructans, fungal carbohydrates and dextran)) ^28^, whereas hydrogenases were classified according to their metal site ^29^. A network based on Spearman’s correlation between the CAZy family and hydrogenases was constructed (Fig. 4a). Network analysis showed that a specific hydrogenase involved in hydrogenotrophy, [FeFe] group A3, was associated with eight different CAZy families involved in the metabolism of animal carbohydrates (mucin- or meat-derived) (Fig. 4a). One plant-based CAZy family, a carbohydrate-binding module known to target the terminal fructoside residue of fructans, was also associated with [FeFe] A3 hydrogenase. Hydrogenase from the [FeFe] group B was associated with seven plant-based CAZy families, including enzymes involved in starch and xylose metabolism. This suggests that the abundance of [FeFe] hydrogenase in gut metagenomes is associated with the metabolism of dietary and host glycans. Finally, we assessed the relationship between the relative abundance of [FeFe] hydrogenases and IBS symptom severity. A linear discriminant analysis showed that [FeFe] A3 hydrogenase was a strong predictor of IBS symptom severity (Fig. 4b). Indeed, [FeFe] A3 hydrogenase presented higher relative abundance in patients with severe IBS compared to healthy subjects (Fig. 4c).

**Figure 4:**
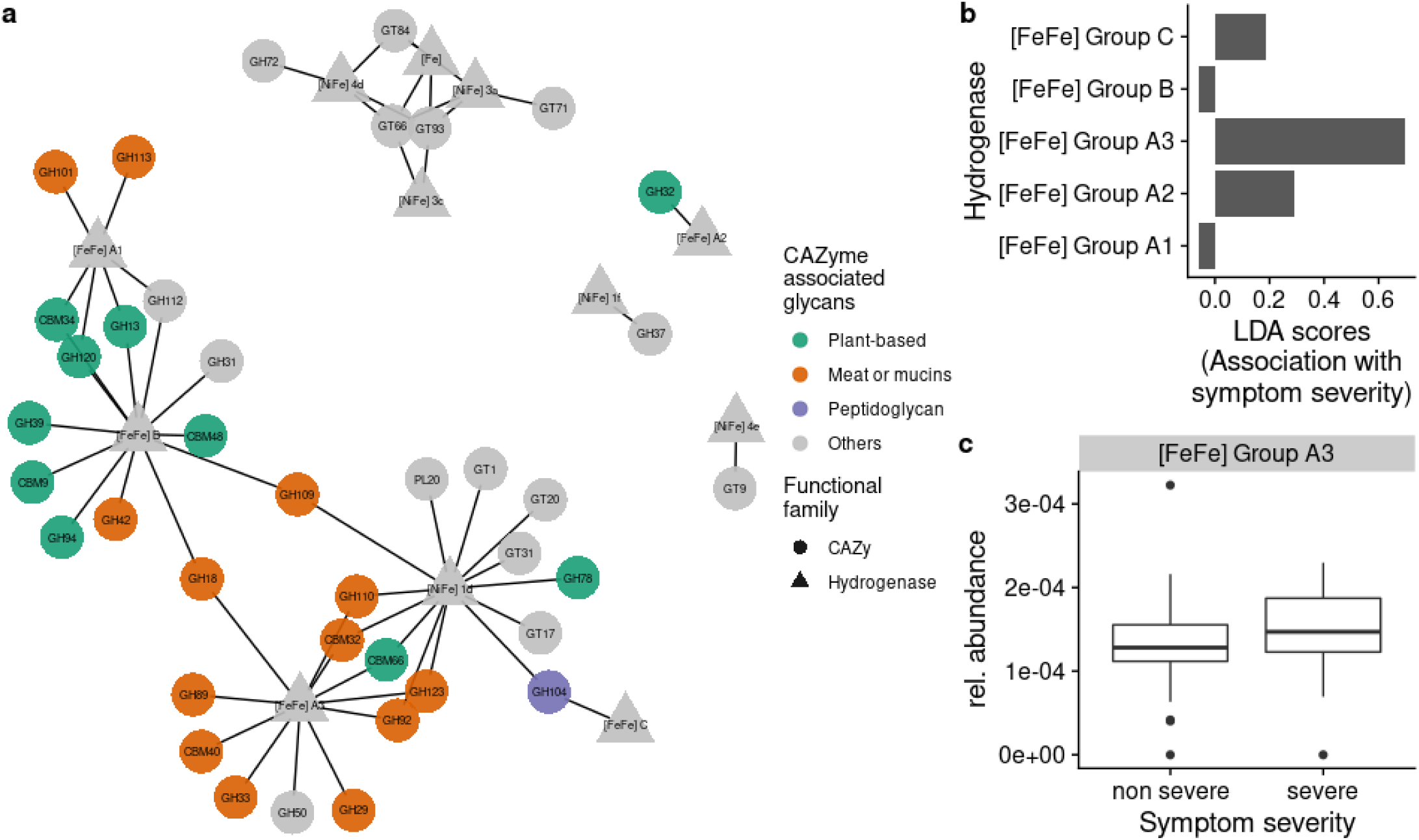
Association between hydrogen levels, glycan metabolism and GI symptom severity. **a** Network of associations between CAZy family and hydrogenase group. An edge represents for absolute Spearman’s correlation coefficients above 0.4. All kept associations were positive. Node shape indicates CAZy family (circle) and hydrogenase (triangle). The color code indicates the known glycan substrates of the CAZy family. **b** Linear discriminant analysis (LDA) scores for prediction of IBS severe vs control as a function of gut metagenomic hydrogenase [FeFe] group. **c** Relative abundance of gut metagenome hydrogenase [FeFe] A3 as a function of health status.

## Discussion

In this study, we showed that patients with severe IBS are characterized by higher intake of poorer quality of food items. Our study further provides evidence that IBS severity is associated with altered gut microbiota hydrogen function in correlation with CAZy involved in animal carbohydrate metabolism. A combination of co-inertia analysis and linear regression analysis suggested that co-variations between gut microbiota and diet could be explained with IBS symptom severity, exhaled gas and glycan metabolism and meat/plant ratio.

Dietary components are often analyzed separately without consideration of their proximity in terms of nutritional value. However, the recently developed approach to structure food items into a hierarchical tree ^4, 14^, similar to that used for microbiota phylogenetic tree analysis, may facilitate the detection of associations that are otherwise difficult to identify, e.g. for nutrients that are not well captured, particularly those known to modulate the microbiota, including fiber and polyphenols ^4^. We took advantage of this approach to decipher diet-gut microbiota interactions in the context of IBS. We modified it, using nutrient intake derived from a 4-day food diary rather than food items themselves, making it possible to separate closely related food items differing by only a few nutrients. Based on the food item tree analysis, meat/plant ratio was the variable that most strongly discriminated subjects from the study cohort. Further, based on FSA-NSP nutrient profiling ^30^, we have shown that the overall quality of dietary intake, of severe IBS patients did not significantly differ from that of other subjects. However, severe IBS patients consumed a higher proportion of lower-quality food items during their main meals. This suggests that quality of food, and not only global diet, is relevant when considering symptoms.

Based on the current finding that the patients with severe IBS symptoms differed from the other subjects in terms of dietary habits, together with our previous identification of a specific gut microbial signature associated with severe IBS symptoms ^23^, we further explored the association between dietary habits and microbiota composition and function Using a new approach combining microbiota and food trees, we identified taxa that were most associated with dietary variation. Analyses of the gut microbiota beyond the species level revealed associations that were not detected in analyses at the species level, and for example *Eubacterium rectale* subspecies harboring flagellin-encoding genes were associated with a predominantly meat-based diet. This complements a previous study that reported that two of the three *E. rectale* subspecies consistently harbored an accessory pro-inflammatory flagellum operon associated with lower gut microbiota community diversity, higher host BMI, and higher fasting blood insulin levels ^31^. These findings highlight the importance of analyzing the gut microbiota at subspecies level, to decipher diet-microbiota-symptom interactions.

We further explored the possible relationship between functional variation in the gut microbiota with dietary and clinical parameters, particularly symptoms, in patients with IBS and healthy subjects. So far, most IBS studies focused on symptoms generated by specific dietary ingredients, and independently of gut microbiota. Using a combination of principal component analysis and linear regression, we were able to predict the diet main variation factor (meat/plant ratio) from gut microbiota profiles of a given subject. Additionally, this approach provides elements to support the hypothesis that dietary profiles and the gut microbiota are respectively associated with IBS symptom severity and exhaled gas resulting from microbial fermentation from dietary ingredients.

As carbohydrates are among the food items that may trigger gas accumulation and symptoms, we further explored the relation between Carbohydrate-associated enzymes (CAZy) and hydrogenases. Herein hydrogenase associated with IBS symptom severity was correlated with higher abundance of CAZy involved in dietary and host animal glycan metabolism. In addition to carbohydrate metabolism, gases, including hydrogen, are of considerable interest in the context of gut disorders ^32^. Hydrogen (H_2_) is formed in large volumes in the colon as an end-product of carbohydrate fermentation ^33, 34, 35^. Recent metagenomics studies have identified three types of hydrogenases involved in H_2_ metabolism ^29, 36^. Interestingly, we found that patients with severe IBS symptoms had a higher abundance of hydrogenases involved in the metabolism of H_2_. This is in agreement with our previous finding that IBS patients with high exhaled H_2_ levels had a distinctive ratio of active members of the gut microbiota regardless of gastrointestinal symptoms ^37^. We confirm the important role of hydrogen metabolism and provide novel insight into the identification of specific hydrogenases for symptom generation in IBS. We suggest that hydrogenases analysis should be encouraged in future IBS and overall diet-microbiota studies.

This study provides a detailed and novel approach that combine both diet and microbiota, but one of its limitations is the lack of longitudinal data and of confirmation in larger, ideally population-based rather than clinical, cohorts. Current dietary records lack resolution into Microbiota Accessible Carbohydrate (MAC), and future microbiota-diet studies would gain in optimising their capture. Nevertheless, this study expands our knowledge of microbiota-diet association and provides new insight into the altered function of the gut microbiota in patients with severe IBS symptoms, potentially as a consequence of interactions with dietary habits. We specifically show that patients with severe IBS symptoms have a higher consumption of lower quality food products and a higher prevalence of microbiota function towards a specific type of hydrogen metabolism associated to animal carbohydrate metabolism. Our findings pave the way for the identification of microbiome-based nutritional recommendations for the management of gastrointestinal symptoms.

## Materials and Methods

### Subject recruitment and study design

Adult patients aged 18-65 years fulfilling the Rome III criteria for IBS ^38^ were prospectively included at a secondary/tertiary care outpatient clinic (Sahlgrenska University Hospital, Sweden). The diagnosis was based on a typical clinical presentation and additional investigations, if considered necessary by the gastroenterologist (HT or MS). Exclusion criteria included the use of probiotics or antibiotics during the study period or within the month preceding inclusion, another diagnosis that could explain the GI symptoms, severe psychiatric disease as the dominant clinical problem, other severe diseases, and a history of drug or alcohol abuse. The healthy control group was recruited through advertisements and checked by interview and with a questionnaire to exclude chronic diseases and any current GI symptoms.

All participants gave written informed consent for participation after receiving verbal and written information about the study. The Regional Ethical Review Board at the University of Gothenburg approved the study before the start of the inclusion period.

### Subject characterization

Demographic information and body mass index were obtained for all subjects. IBS patients reported their current use of medications and completed questionnaires to characterize their symptom severity and bowel habits: The IBS Severity Scoring System (IBS-SSS)^26^, a 4-day food diary and a two-week stool diary based on the Bristol stool form scale. IBS severity was assessed with validated cutoff scores for the IBS-SSS (mild IBS: IBS-SSS <175, moderate IBS: IBS-SSS=175-300, severe IBS: IBS-SSS>300). Oral-anal transit time (OATT) (radiopaque marker study)^39^ and exhaled H_2_ and CH_4_ levels after an overnight fast (i.e. with no substrate intake preceding the test) were also determined for IBS patients (see supplementary online-only material for more details). Exhaled CH_4_ and H_2_ levels were determined after an overnight fast (i.e., not after the intake of any substrate), and after the subjects had received thorough instructions to avoid a diet rich in fiber and poorly absorbed carbohydrates the day before the test. Exhaled H_2_ and CH_4_ levels were determined in parts per million in end-expiratory breath samples collected in a system used for the sampling and storage of alveolar air (GaSampler System; QuinTron Instrument Company, Milwaukee, WI) immediately analyzed in a gas chromatograph (QuinTron Breath Tracker; QuinTron Instrument Company).

### Dietary intake

All subjects completed a paper-based diet record, in which all foods and drinks consumed during four consecutive days (Wednesday-Saturday) were reported. Oral and written instructions were given to the patients on how to record their dietary intake, and patients were told to keep to their regular diet during the recording days. The type of food and the time at which it was consumed were noted, with quantification in grams, according to the use of household utensils (e.g. tablespoons) or number of slices, for example. Cooking method and the contents of food labels were noted where applicable. All diet records were entered into a special version of Dietist XP 3.1 software (Kostdata.se, Stockholm, Sweden), which calculates the energy and nutrient composition of foods. The software was linked to a Swedish Food Composition Database provided by the National Food Agency in Sweden (https://www.livsmedelsverket.se/), and to a Swedish database with FODMAP content, developed in-house ^40^. This database contained information about fructose, fructan, lactose, galacto-oligosaccharide (GOS) and polyol content (g/100g) from published sources (Storsud et al, submitted). All diet records were entered into the software by a trained dietician. Excess fructose levels were calculated from data for fructose and total monosaccharide content from diet records. Glucose and fructose are the dominant monosaccharides in foods. If glucose content was higher than fructose, then the excess fructose variable was assigned a value of 0 (for each separate meal). Nutrient intakes were first summarized for each meal, and then per day, and were finally determined as mean intake for all four days. A cutoff value was set for energy intake, and subjects reporting energy intake levels below 800 kcal/day or exceeding 4500 kcal/day were excluded, to remove reports corresponding to an implausible habitual intake. No subjects exceeded these limits. For the reported intake of FODMAPs, outliers were defined as values exceeding the mean±4 SD.

### FSA-NPS diet index

The FSA-NPS (British Food Standards Agency Nutrient Profiling System) ^41^ score was calculated for all foods and beverages, as follows: points (0–10) are allocated for the content per 100 g in total sugars (g), saturated fatty acids (g), sodium (mg), and energy (kJ) and can be balanced by opposite points (0–5) allocated for dietary fiber (g), proteins (g), and fruits/vegetables/legumes/nuts (percent). The grids for point attribution were as described by Deschasaux et al. ^30^. The FSA-NPS score for each food/beverage is based on a unique discrete continuous scale ranging theoretically from −15 (most healthy) to +40 (least healthy). In addition, each food item was assigned to one of five groups: from A (high quality), to E (low quality) for a FSA-NPS score below 1, 2, 5, 9, 40 for A, B, C, D and E, respectively, for drinks, and below 0, 3, 10,18 and 40 for A, B, C, D and E, respectively, for all other foods. The overall diet index, FSA-NPS DI, was calculated as the energy-weighted mean of the FSA-NPS scores of all foods and beverages consumed, as described by Deschasaux et al. ^30^.

### Hierarchical food tree and UniFrac analysis

We used the hierarchical format of the Swedish Food Composition Database to categorize foods into a hierarchical tree, the food tree. Food items, their associated nutrients and their corresponding hierarchical levels were downloaded from the Swedish Food Composition Database (https://www.livsmedelsverket.se/). These hierarchical levels corresponded to levels 3 and 4 in the food tree. We then grouped level 2 into five large categories: animal-based, plant-based, alcohol, fats and others. Level 1 is the root of the food tree. We then divided level 4 into 85 subcategories, constituting level 5 in the food tree, as follows:

1. For each level 4 category, we extracted the nutrient content for each food
2. We fitted a Dirichlet multinomial mixture (DMM) model to each category.
3. The number of Dirichlet components that resulted in the minimum Laplace approximation have been selected
4. Each DMM component was used to assign food items to a subcategory, in level 5 of the food tree.

The hierarchical structure of the food tree is shown in Supplementary Table S1. We used the food tree to calculate unweighted UniFrac metrics between food diaries.

### Fecal sample collection and DNA extraction

Fecal samples from 138 subjects were collected in RNA Later solution (Ambion, Courtaboeuf, France). Fecal DNA was extracted by mechanical lysis (Fastprep® FP120 (ThermoSavant)) followed by phenol/chloroform-based extraction, as previously described ^23^. A barcoded fragment library was prepared for each sample, and DNA sequencing data were generated with SOLiD 5500xl sequencers (Life Technologies), resulting in a mean of 38 (SD 14) million sequences of 35-base single-end reads. High-quality reads were generated, with a quality score cutoff > 20. Reads with a positive match to human, plant, cow or SOLiD adaptor sequences were removed. Filtered high-quality reads were mapped onto the MetaHIT 3.9 million genes catalogue with METEOR software. The read alignments were performed in color space with Bowtie software (version 1.1.0), with a maximum of three mismatches and selection of the best hit. Uniquely mapped reads (reads mapping to a single gene from the catalogue) were attributed to the corresponding genes and used to construct a raw gene count matrix. If multiple alignments were found, counts were divided equally between the aligned genes.

### Metagenomics species pangenome analysis

Metagenomics species pangenomes (MSPs) are co-abundant gene groups that can be considered part of complete microbial species pangenomes. MSP gene content was extracted from a previous publication by Plaza-Onate et al. ^27^. MSP gene content was subdivided into core and accessory genes. Gene annotations (KEGG orthology and CAZy family) were extracted from the paper by Li et al. ^42^. Thus, MSP relative abundance was calculated for each sample, based on median core gene abundance. Samples were attributed to an MSP subspecies on the basis of accessory gene clustering, as follows:

1. Median read coverage and the 2.5 and 97.5% quantiles were calculated for each sample and MSP. An MSP was considered to be detected in the sample if it had a median coverage of more than 2.
2. Each accessory gene within the MSP with a read coverage between the 2.5 and 97.5% quantiles was considered to be detected. Below the 2.5% quantile, genes were considered to be absent, and above the 97.5% quantile, genes were considered to be present in multiple copies or to be conserved genes that might bias the estimation of coverage. A presence/absence binary gene matrix was therefore obtained for each MSP.
3. A Jaccard index between samples was calculated from the MSP binary matrix
4. Clustering was performed, with a partition around medoids over 100 bootstraps achieved with the *clusterboot* function of the *fpc* package (bootstrap method option *“subset”*). The number of clusters (i.e. MSP subspecies) was estimated on the basis of mean silhouette width.

### Gut microbiome hydrogenase and CAZy analysis

Hydrogenase amino-acid sequences were extracted from a previous study ^29^ and aligned, with BlastX software (version 2.7.1+), with 3.9 million gene catalogs. Best hits with an identity of more than 60% over a stretch of more than 40 amino acids were considered for downstream analysis.

### Statistical analysis

All statistical analyses were performed with R software (version 3.4.1). UniFrac distances between food diaries were calculated with the phyloseq R package, using the food tree as input. JSD distances between metagenomes aggregated at MSP subspecies level were calculated according to a tutorial published by Arumugam et al.^43^ implemented into BiotypeR R package (available on github tapj/BiotypeR). PERMANOVA analysis was performed with the vegan R package (Adonis, version 2.5). Principal coordinate analysis (PCoA), co-inertia analysis and linear discriminant analysis were performed with the ade4 R package (version 1.5). To note, PCoAs on microbiota and diet distance were computed using all available samples while co-inertia analysis was computed on common sub-samples from PCoAs components. Spearman correlation analysis was used to project features onto PCoA axes. Principal component linear regression analysis was used to train models to predict clinical, microbiome and diet features (e.g. exhaled gas metabolism, symptom severity, meat/plant ratio) with the co-inertia axes. Spearman’s correlation analysis was performed on relative abundance data for genes aggregated at the CAZy family and hydrogenase levels. A network was generated for correlations with an absolute rho value above 0.4. Data were visualized with cowplot, ggraph and ggplot2. *P* values were adjusted for multiple testing by Benjamini–Hochberg false-discovery rate correction when specified. Otherwise, p.values are given at 5% nominal level.

## Specific author contributions

Designed the study: HT, LO, MS; conducted the study/collected data: HT, LO, SS, MS; were responsible for sequencing and sample handling: NP, JD; analysed the data: JT, SS; interpreted the study: AC, JT, BLN, MD, MS. Drafted the manuscript: MD, JT; Commented on the drafts of the paper: all co-authors; approved the final draft submitted: all co-authors

## Availability of data and materials

The datasets used and analyzed in this and the parent study ^23^ are available from: https://doi.org/10.5878/ejpj-p674, https://doi.org/10.5878/5rbz-ww62.

## Acknowledgments

The authors wish to thank Stéphanie Cools-Portier and Sean Kennedy for logistical or analytical support with metagenomic sequencing, Heleen de Weerd, Martin Balvers and Jolanda Lambert for bioinformatics support, Patrick Veiga for critical reading of the manuscript.

## Supporting information

Supporting Materials and Methods

Supporting Figures

Supporting Tables

## Supporting Figures

**Figure S1.**
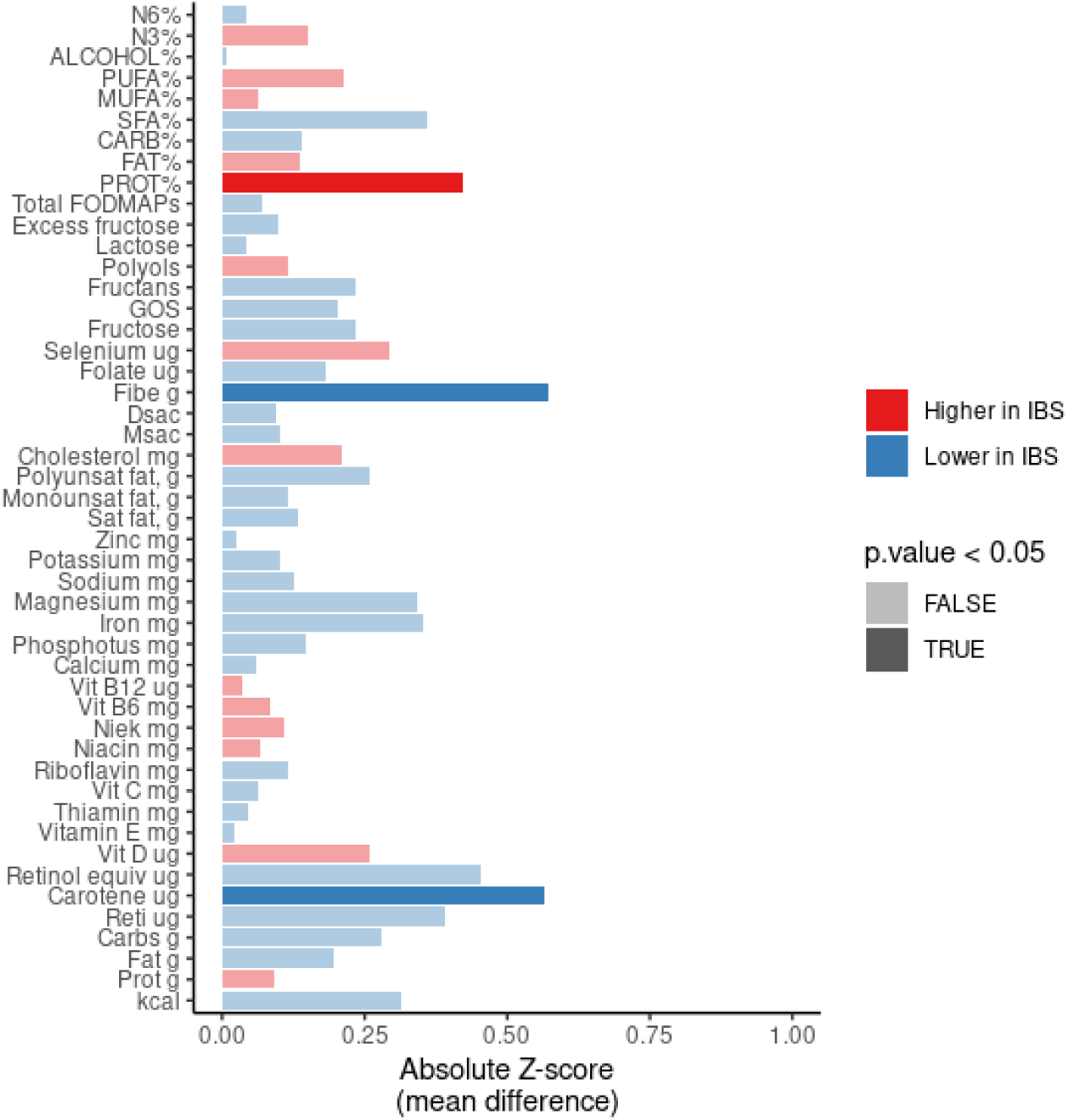
Nutrients more abundant in the diets of healthy controls than in those of IBS patients. Absolute *Z*-scores between controls and IBS patients are shown. A positive *Z*-score indicates an enrichment of the diet in the nutrient concerned and, is shown in red for healthy subjects and in blue for IBS patients. Significant *p*-values (uncorrected for multiple tests) are shown in a darker color.

**Figure S2.**
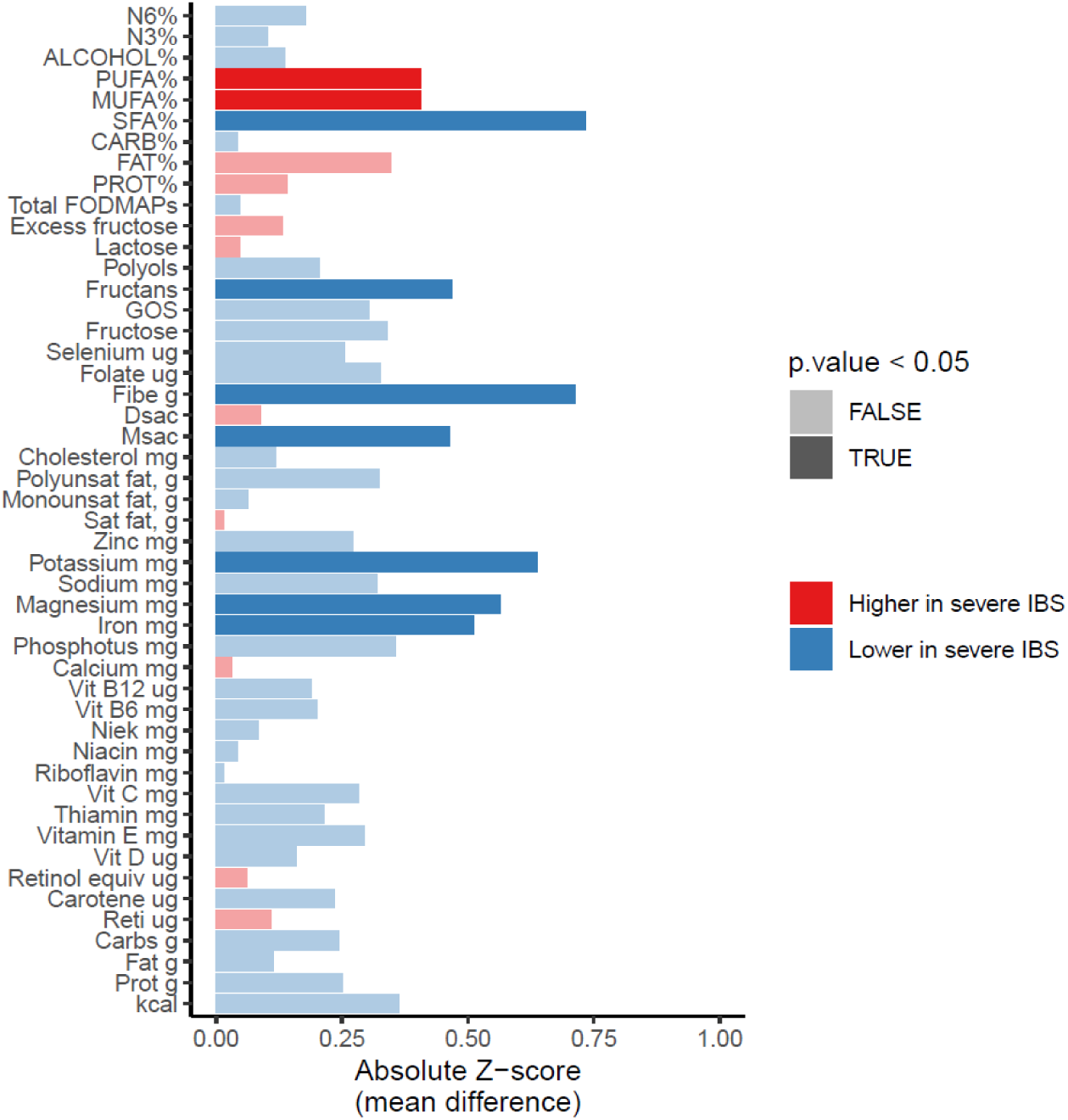
Comparison of nutrients between IBS patients with severe symptoms and the other members of the study population (healthy, or with mild or moderate IBS). Absolute *Z*-scores for comparisons between IBS patients with severe symptoms and the other members of the study population are shown. Positive *Z*-scores, indicating nutrient depletion in IBS patients with severe symptoms are shown in blue. Significant *p*-values (uncorrected for multiple testing) are shown in a darker color.

**Figure S3.**
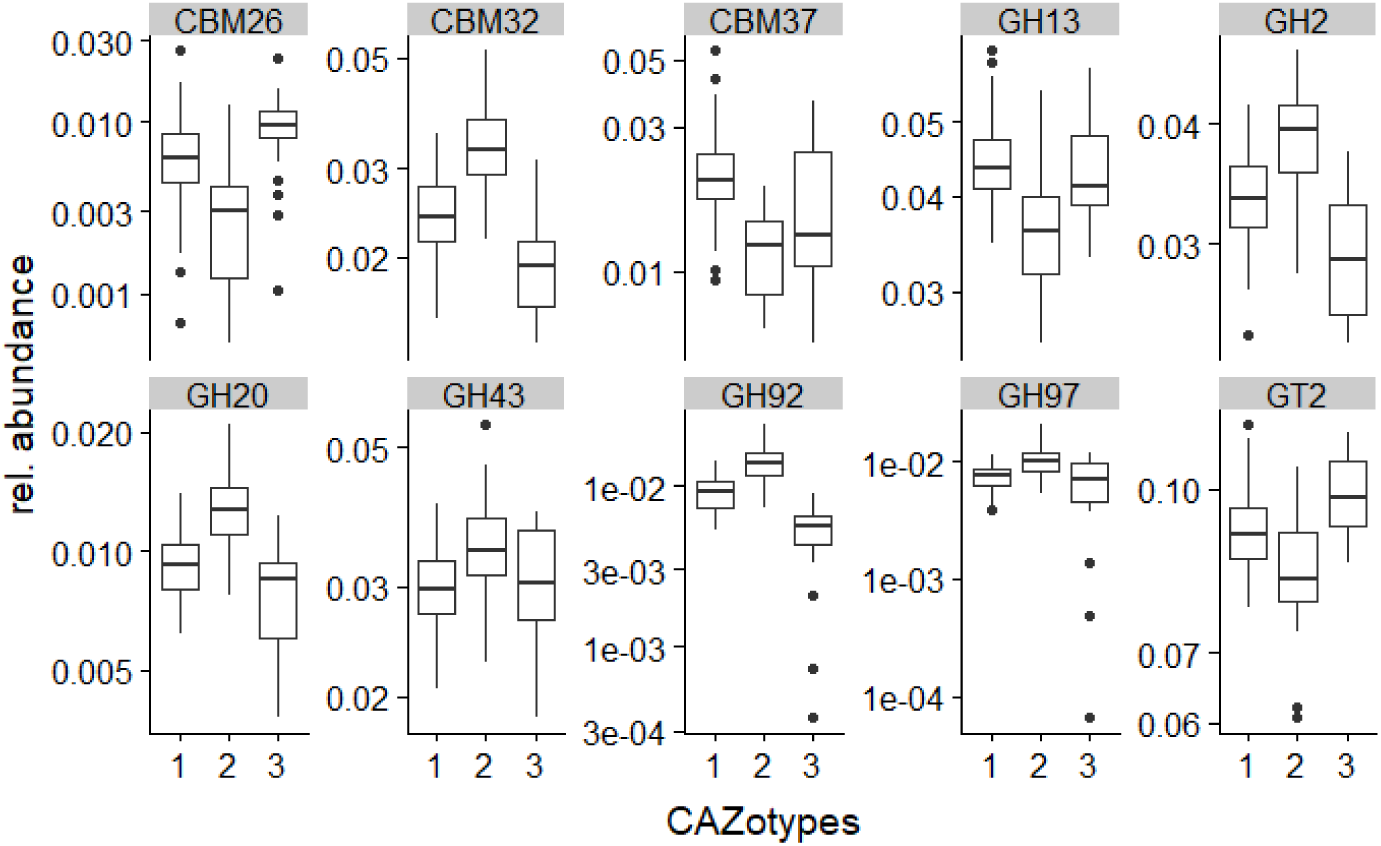
CAZotype as a function of the relative abundance of CAZy genes. CAZotypes 1 and 3 were notably enriched in glycosyl hydrolase (GH) 13 and carbohydrate binding module (CBM) 26, both involved in starch metabolism, whereas CAZotype 2 was enriched in GH2 and CBM32. CAZotype 3 displayed a particular depletion of GH2 and GH20, which are known to be involved in mucin degradation.

**Figure S4.**
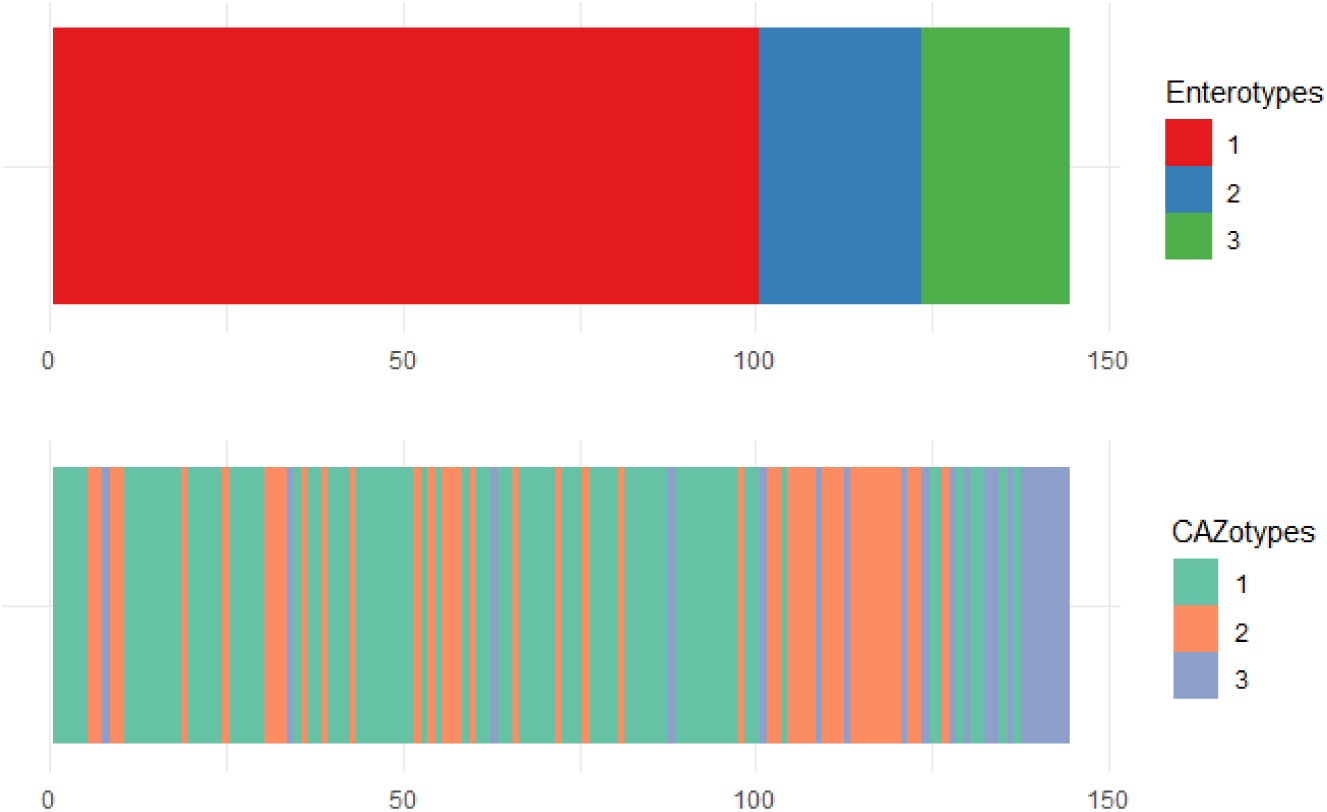
CAZotype and enterotype assignment, by individual.

## Supporting Tables

**Table S1.** Food items studied and their associated hierarchical classification

**Table S2:** meat plan ratio and diet based PCoA components per individual

**Table S3:** Spearman correlation analysis between clinical and microbiome variable with co-inertia components

